# Stable heteroplasmy at the single cell level is facilitated by inter-cellular exchange of mtDNA

**DOI:** 10.1101/007005

**Authors:** Anitha D Jayaprakash, Erica Benson, Swapna Gone, Raymond Liang, Jaehee Shim, Luca Lambertini, Masoud M Toloue, Mike Wigler, Stuart A Aaronson, Ravi Sachidanandam

## Abstract

Eukaryotic cells carry two genomes, nuclear (nDNA) and mitochondrial (mtDNA), which are ostensibly decoupled in their replication, segregation and inheritance. It is increasingly appreciated that heteroplasmy, the occurrence of multiple mtDNA haplotypes in a cell, plays an important biological role, but its features are not well understood. Accurately determining the diversity of mtDNA has been difficult, due to the relatively small amount of mtDNA in each cell (< 1% of the total DNA), the intercellular variability of mtDNA content and mtDNA pseudogenes (Numts) in nDNA. To understand the nature of heteroplasmy, we developed Mseek, a novel technique to purify and sequence mtDNA. Mseek yields high purity (> 90%) mtDNA and its ability to detect rare variants is limited only by sequencing depth, providing unprecedented sensitivity and specificity. Using Mseek, we confirmed the ubiquity of heteroplasmy by analyzing mtDNA from a diverse set of cell lines and human samples. Applying Mseek to colonies derived from single cells, we find heteroplasmy is stably maintained in individual daughter cells over multiple cell divisions. We hypothesized that the stability of heteroplasmy could be facilitated by inter-cellular exchange of mtDNA. We explicitly demonstrate this exchange by co-culturing cell lines with distinct mtDNA haplotypes. Our results shed new light on the maintenance of heteroplasmy and provide a novel platform to investigate features of heteroplasmy in normal and diseased states.

## Introduction

Mitochondria are organelles present in almost every eukaryotic cell [1]. They enable aerobic respiration[2] to efficiently generate ATP, and play an important role in oxygen sensing, inflammation, autophagy, and apoptosis[3, 4]. Mitochondrial activity relies on over a thousand proteins, mostly coded by the nuclear DNA in humans[5], but proteins from the mitochondrial genome, a small circular DNA (**mtDNA**), play a critical role in their function. In humans, the reference mtDNA is 16,569 bp long and codes thirteen proteins critical for the electron transport chain, along with twenty-two tRNAs, two rRNAs and a control region, called the displacement loop (D-loop) (Fig. S1)[6]. Each mitochondrion carries multiple mitochondrial genomes (5 –10)[7] and each cell contains hundreds to thousands of mitochondria, depending on the tissue[8]. The mtDNA replicate without recombination. mtDNA is inherited solely from the mother; inherited mutations in mtDNA have been linked to several genetic disorders including *diabetes mellitus and deafness(DAD)* and *Leber’s hered-itary optic* neuropathy(LHON)[9]. De novo mutations in mtDNA have also been linked to diseases[10, 11, 12, 13].

Heteroplasmy, which is the occurrence of multiple mtDNA haplotypes, has been documented in various studies, cancer cells[14, 15], blood samples from families[16] blood and muscle biopsies from identical twins[17], and cells from the 1000 genomes project [18, 19]. Though extensive, these studies have not established the nature of heteroplasmy at the *level of the single cell*. Accurate determination of heteroplasmy, especially the low-frequency haplotypes, is needed for disease-association studies with mtDNA, as well as studies of metabolic activity of cancer cells[14, 20]. Deep sequencing is the only means to identify novel mtDNA haplotypes as well as somatic mutations in tissues and perform association studies to link the haplotypes to disease states. However, measurements of heteroplasmy are compromised by copies of large segments of mtDNA, called nuclear mitochondrial DNA (**Numts**), present in the mammalian nDNA[21] (Fig. S2, S3).

Without purification of mtDNA, Numts introduce unpredictable inaccuracies in the estimates of heteroplasmy, especially because they exhibit variations in sequence and copy numbers. Numts have been annotated in the reference human genome[22, 23], and there are tools to analyze high throughput sequencing data in light of these annotations [24], but a comparison of two recent versions of the reference human genome (hg19 and hg38 on the UCSC genome browser, Fig. S2, S3) suggests that this annotation is not complete and changes significantly with the reference genome. Additionally, the distribution of Numts might be specific to each individual’s nuclear genome. A recent study of mtDNA from twins has highlighted the need for further investigation of Numts and their potential to confound analyses of heteroplasmy[25].

Isolating mtDNA has long been a challenge. In forensics and genealogy, allele-specific primer extensions (SNaPshot) are used for genotyping mtDNA[26]. Hyper variable regions (HVR) in the D-loop have been amplified using PCR[27]. Entire mtDNA has been accessed using primers specific to mtDNA to either perform long-range PCR[28], or amplify overlapping fragments[14]. Isolation of organelles by ultra-high-speed centrifugation[29] has also been used, but the yields are low and contaminated with fragmented nuclear DNA[30]. Computational methods have also been used to infer heteroplasmy from whole-exome[19, 31] and whole-genome data[15], but such data contains Numts and furthermore, generating such data for new samples is expensive. A new approach uses methyl-specific endonucleases MspJI and AbaSI to deplete nDNA that is likely to be methylated[32], but Numts need not be methylated. Heteroplasmy derived from PCR-based methods are error-prone, due to variability in amplification. Clonal amplification of errors introduced by polymerases is also a problem in PCR-amplicon sequencing. Additionally, sequence and copy number variations of Numts confound results from computational and PCR-based methods in unpredictable ways. Thus, it is difficult to ascertain if the methods outlined above are accurate in their measurement of frequencies of common variants and their identification of variants that occur at frequencies below 5%.

We present here Mseek, a novel method to enzymatically purify mtDNA by depleting linear nDNA and inexpensively sequencing it (Fig. S4). Mseek uses exonuclease V to digest away linear nDNA, leaving behind circular mtDNA. A major benefit of this method is the ability to call extremely rare variants, with the sensitivity of calls only limited by the sequencing depth. By applying Mseek to several cell-lines and human peripheral blood mononuclear cells (PBMC), we identified mixtures of different mtDNA haplotypes in the samples. Additionally, through clonal expansion of single cells from a variety of cell lines, we establish, for the first time, that heteroplasmy is stably maintained at a single cell level over multiple divisions. We infer the stability is due to intercellular exchange of mtDNA and experimentally demonstrate this exchange by co-culturing two different cell-lines with distinct mtDNA haplotypes to show that there is transfer of mtDNA from one cell type to another.

## Results

### Mseek: An Efficient Method to Isolate and Sequence mtDNA

To efficiently purify mtDNA, we sought to take advantage of the difference in topology between nDNA and mtDNA using an exonuclease to digest the linear nDNA, while leaving intact the circular mtDNA. Total DNA was extracted from HEK 293T cells, and digested with exonuclease V or left undigested. To determine the efficiency of digestion, sequences specific to nDNA and mtDNA were PCR amplified using appropriate primers (Tables ST1 and ST2). As expected, in the undigested samples of total DNA we detected both nDNA and mtDNA (Fig. 1A). In sharp contrast, in the samples treated with exonuclease V we only detected mtDNA (Fig. 1B). The lengths of the expected PCR products are shown in Fig. 1C.

**Figure 1.**
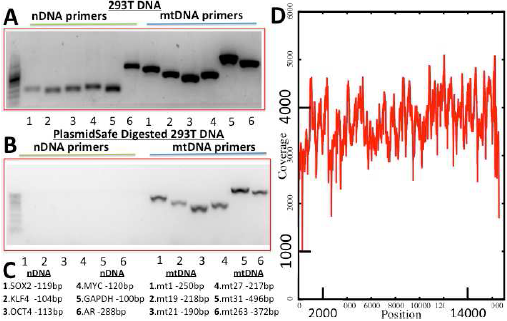
Performance of Mseek. (A) PCR products run on a 2% Agarose gel using primers for 6 nDNA genes (OCT4, MUC, KLF4, SOX2, GAPDH and AR, Table ST1) and 5 regions of mtDNA (Table ST2) before exonuclease digestion. (B) After digestion, the nDNA bands disappear. (C) Sizes of expected PCR products. (D) Deep sequencing, read depth (y-axis) versus position on mtDNA (x-axis) demonstrating uniformity of coverage. 1.23 million mtDNA reads and 50,000 nDNA reads implying > 98% pure mtDNA.

Using this approach, mtDNA was prepared and sequenced on the Illumina MiSeq platform. Out of a total of 3.05 million 100nt reads, 1.233 million mapped to the mitochondrial genome and 50,000 (< 2%) mapped to the nDNA. The rest of the reads were adapter dimers, formed during library preparation by the ligation of adapters to each other. Over 98% of the mappable reads were derived from mtDNA with an average coverage > 3000X (Fig. 1D). More than 50 distinct samples were processed similarly to consistently obtain high purity mtDNA. This approach, designated **Mseek** (Fig. S4), provides a means of unmatched efficiency in accurately sequencing the mtDNA contained within a population of cells.

Our comparisons of Mseek to other kits on the market show that Mseek substantially outperforms all of them in terms of the purity of mtDNA and yield (Supplementary material and Fig. S5). As currently implemented, a limitation of Mseek is its requirement of at least 4 *μ*grams of intact total DNA, which is a big improvement over the 50*μ*g of total DNA that were needed for the initial experiments. Further improvements of sample preparation may reduce the input amount required, but are beyond the scope of this paper. For less than 4*μ*g of total DNA we recommend using long-range PCR amplification with mtDNA specific primers after exonuclease V treatment, which depletes nDNA to minimize distortions arising from Numts. The primers for long range PCR are specified in table ST2 and enzyme used for amplification is from Clontech (Advantage Genomic LA Polymerase Mix catalog # S4775).

Multiple rounds of treatment with exonuclease V maybe needed to achieve high purity, a single round usually achieves around 80% purity. Assuming around 20 copies of the mtDNA per nuclear genome (which is probably an overestimate), the contamination from Numts is less than 0.01%, which can be safely ignored for most applications. Thus, Mseek can be used to call rare variants to any level of sensitivity, only limited by the depth of sequencing. Most methods will not allow this, for example, in PCR-amplicon sequencing, sensitivity does not necessarily increase with depth of sequencing as errors introduced during amplification cannot be corrected by greater sequencing depth. This is a valuable feature, that distinguishes it from other techniques, enabling the tracking of rare variants. Duplex sequencing, which uses adaptors with random tags to track clones in order to reduce the errors introduced during sample preparation for deep-sequencing, should be used in conjunction with Mseek to identify and study rare variants (with frequencies < 0.1%) [33].

### Ubiquity of Heteroplasmy

Since cell lines are clonally derived, the nDNA (and mtDNA) are expected to be identical across cells, since, either i) a slight fitness advantage of one haplotype or ii) drift[34], would lead to clonal selection and homoplasmy. To study clonality in mtDNA, we applied Mseek to thirty samples including four human PBMCs and human cell lines derived from human diploid fibroblasts (501T), glioma (A382) and breast carcinoma (HCC1806 and MDA-MB-157). Repeat content of the sequences was computationally identified to estimate nDNA contamination, which ranged from 0.5 − 5%; confirming the specificity of Mseek. Importantly, because of this high degree of mtDNA purity, we were able to multiplex all 30 samples in a single MiSeq run, with average coverage of > 50X. We show the coverage at various positions in the Tables 1 and 2.

**Table 1.**
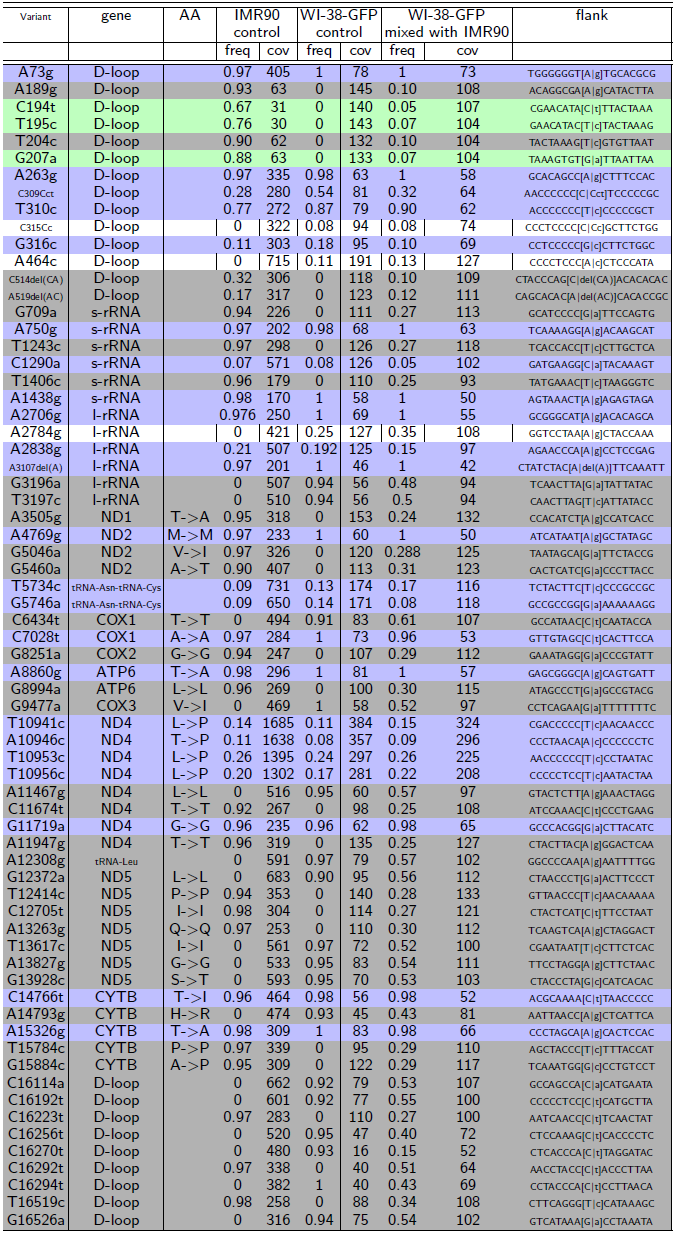
Co-culturing experiments for WI-38-GFP(fibroblast) and IMR90 (fibroblast). The white rows are variants private to WI-38-GFP, gray rows show variants transferred from IMR90 to WI-38 and blue rows are variants common to IMR90 and WI-38. The green rows are variants private to IMR90 with minimal or no transfer. The frequency (*freq*) ranges from 0-1 and the coverage (*cov*) is the number of reads at the variant.

**Table 2.** Data from mixing experiments of MDA-MB-157 cells co-cultured with HCC1806 cells for 6 weeks. The MDA-MB-157 cells were labeled with GFP and made up approximately 10% of the mixture, which was passaged approximately 25 times before the component cells were separated using FACS. The mtDNA was sequenced to identify variants in the GFP and non-GFP cells. The private_to column identifies the cell-lines that exhibit the variant. Rows highlighted in gray are cases where a variant unique to HCC1806 has been identified in MDA-MB-157 cells. The light green rows are variants private to HCC1806 that did not transfer into MDA-MB-157. Rows highlighted in blue show variants common to both cell lines. For example, at position 3796 (row 6), the A from the reference mtDNA genome is mutated to a *T* only in HCC1806, the MDA-MB-157 cultured with HCC1806 exhibits an *A*, with a frequency of 0.14 ( or 14%). The frequency is *freq* (which ranges from 0-1) and the coverage, number of reads covering the variant, is *cov*.

| variant | gene | AA | private_to | HCC1806 control |  | MDA-MB-157 from HCC1806 |  | flank |
| --- | --- | --- | --- | --- | --- | --- | --- | --- |
|  |  |  |  | freq | cov | freq | cov |  |
| G3666A | ND1 | G120G | HCC1806 | 0.93 | 33 | 0 | 25 | TCTGATCAGG[G\|a]TGAGCATCAA |
| A9545G | COX3 | G113G | HCC1806 | 0.93 | 15 | 0 | 13 | CCCAATTAGG[A\|g]GGGCACTGGC |
| A12810G | ND5 | W158W | HCC1806 | 0.92 | 25 | 0 | 20 | AATCCTATAC[A\|g]ACCGTATCGG |
| C14911T | CYTB | Y55Y | HCC1806 | 0.96 | 29 | 0 | 39 | CTATATTACG[C\|t]ATCATTCTCTC |
| C16187T | D-loop |  | HCC1806 | 1 | 9 | 0 | 31 | ATCAAAACCC[C\|t]CTCCCATGC |
| A3796T | ND1 | T163S | HCC1806 | 0.88 | 9 | 0.14 | 21 | TAACCTCTCC[A\|t]CCCTTATCAC |
| A4104G | ND1 | L266L | HCC1806 | 0.9 | 21 | 0.06 | 31 | TTCTAACCT[C\|a]CTGTCTTAT |
| T7861C | COX2 | D92D | HCC1806 | 0.96 | 29 | 0.05 | 39 | CGGACTAAT[C\|t]TCAACTCCTA |
| G9064A | ATP6 | A179T | HCC1806 | 1 | 10 | 0.16 | 24 | CCATTAACCT[G\|a]CCCTCTACAC |
| A9072G | ATP6 | S182S | HCC1806 | 0.88 | 9 | 0.18 | 22 | CTTCCCTCTA[A\|g]ACTTATCATC |
| G10688A | ND4L | V73V | HCC1806 | 0.96 | 26 | 0.1 | 19 | ACCTGACTCC[G\|a]ACCCCTCACA |
| C7819A | COX2 | L78L | MDA-MB-157 | 0 | 35 | 0.76 | 43 | CGCATCCTTT[C\|a]CATAACAGAC |
| C8932T | ATP6 | P135S | MDA-MB-157 | 0 | 25 | 0.9 | 21 | CCATACTAGT[C\|t]ATTATCGAAA |
| T13602C | ND5 | Y422Y | MDA-MB-157 | 0 | 22 | 0.45 | 24 | CAAGCGCCTA[T\|c]AGCACTCGAA |
| T15514C | CYTB | Y256Y | MDA-MB-157 | 0 | 12 | 0.89 | 28 | CAGACAATTA[T\|c]ACCCTAGCCA |
| T16209C | D-loop |  | MDA-MB-157 | 0 | 17 | 0.92 | 26 | TACAAGCAAG[T\|c]ACAGCAATCA |
| C16292T | D-loop |  | MDA-MB-157 | 0 | 15 | 0.76 | 21 | CAAACCTACC[C\|t]ACCCTTAACA |
| C16295T | D-loop |  | MDA-MB-157 | 0 | 14 | 0.72 | 22 | ACCTACCCAC[C\|t]CTTAACAGTA |
| T9540C | COX3 | L111L | Both | 0.86 | 15 | 1 | 11 | TACCCCCCAA[T\|c]TAGGAGGGCA |
| C16223T | D-loop |  | Both | 0.95 | 20 | 0.96 | 31 | GCAATCAACC[C\|t]TCAACTATCA |
| T16311C | D-loop |  | Both | 0.75 | 12 | 0.8 | 20 | CAGTACATAG[T\|c]ACATAAAGCC |

The sequences were analyzed for variants using MiST[35]. Variants with a frequency of 1 indicate homoplasmic mtDNA. Frequencies less than 1 imply the coexistence of multiple haplotypes. Strikingly, in both cell-lines and human blood-derived mtDNA, we observed variants occurring in the 0.1 − 0.9 frequency range (Fig. 2), indicating the presence of multiple haplotypes. Most mutations were transitions (Table ST3), suggesting that the mutations don’t arise from oxidative stress, and are most likely driven by the polymerase-γ activity[33]. The program *Mutation Assessor*[36] was used to label the variants as *high, medium, low*, or *neutral* signifying their predicted impact on protein function. Cell-lines and human PBMCs did not exhibit mutations of putative high effect at a high frequency (> 5 %), consistent with the expectation that functioning cells should have functional mitochondria.

**Figure 2.**
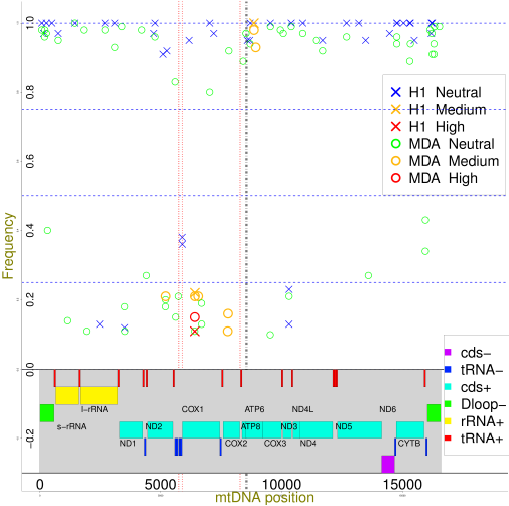
Variants in a cell-line (MDA-MB-157, circles) and a human (PBMC, crosses) compared to the reference mtDNA sequence. Mutation frequency (y-axis) versus position on mtDNA (x-axis). Genes are colored bands at bottom of graph (+, − represent the strand, cds is coding sequence). *Neutral, Low, Medium*, and *High* are the effect of the mutation on protein function (*Mutation Assessor*[36]). Except for the D-loop, most of the mtDNA codes for a transcript, with a few gaps. The longer gaps (11, 24, and 30 nt long) are marked by vertical red lines. The 45 nt long overlapping region between ATP8 and ATP6 is marked by black vertical lines. Mutation frequencies between 0 and 1 arise from the co-existence of multiple mtDNA haplotypes. Heteroplasmy at a cellular level is demonstrated in Fig. 3. There are no stark differences between the human and cell-line derived mtDNA.

Each sample had unique, distinguishing mutations, ranging in frequency from 0.36 to 1.0. There were a number of variants unique to each of the human PBMC samples (ranging in number from 5 to 15) and each of the cell lines (ranging in number from 5 to 21). Our findings hold for cell lines derived from a variety of tissues, suggesting they are of general applicability. There were no key distinguishing features between normal or cancer cell lines and human blood-derived mtDNA, in terms of deleterious mutations or degree of heteroplasmy.

### Stability of Heteroplasmy in Cell-Lines

The results above indicate that heteroplasmy exists within a cell population but questions remain about the nature of heteroplasmy in individual cells. A mixture of homoplasmic cells with different haplotypes can appear to be heteroplasmic. In order to establish heteroplasmy in individual cells, we placed the severest possible bottleneck on the population by deriving colonies from single cells, utilizing MDA-MB-157 and U20S breast carcinoma and osteosarcoma lines respectively (Fig. 3). In each of the derived colonies (4 colonies per cell line), variants from the original lines existed in the derived colonies at approximately the same frequencies as in the original colonies. The sharing of mutations between the original and derived colonies implies the diversity in mtDNA is present in individual cells. The preservation of the frequencies between the original and derived colonies indicates further that this heteroplasmy is uniform across cells in the original line (Fig. 3). Low frequency mutations (frequency < 2%) in the original colony do disappear in the derived colonies, suggesting the rarer variants might be present in a subset of cells, unlike the more common variants, and might be transient.

**Figure 3.**
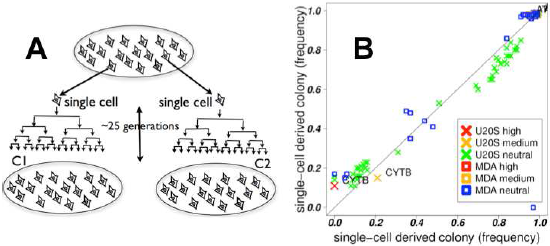
Stability of heteroplasmy. A C1 and C2 are colonies derived from single cells in the original colony, placing a severe botteneck on the mtDNA, and then passaged ≈ 25 times. B Mutation frequencies in C1 (x-axis) versus C2 (y-axis) for two cell-lines, U20S and MDA-MB-157. The mutations mostly lie along the diagonal; the heteroplasmic mix of mutations in the derived colonies are similar to each other. This implies the heteroplasmic mix exists at the single-cell level, and is stable over many divisions. A drift in frequencies is expected with random assortment of mtDNA haplotypes (simulations, Fig. 4). The stability of the frequencies implies active mechanisms to counteract the drift, such as the exchange of mtDNA between cells.

A simple model of mtDNA genetics assumes random assortment of mtDNA haplotypes between daughter cells upon cell division, followed by multiplication of mitochondria. This model would predict drift towards homoplasmy, as seen in our simulation of this process (Fig. 4) and by others[34]. The rate of drift in haplotype frequencies is a function of the number of mtDNA molecules per cell and the original frequency of the haplotypes (Fig. 4). After many passages, irrespective of the original mtDNA distribution, the likelihood of two randomly selected cells having the same heteroplasmic mix would be extremely low, which is at odds with the stable and uniform heteroplasmy that we observed in the clonally-derived cell-lines. This suggests the existence of an active mechanism to counteract this drift.

**Figure 4.**
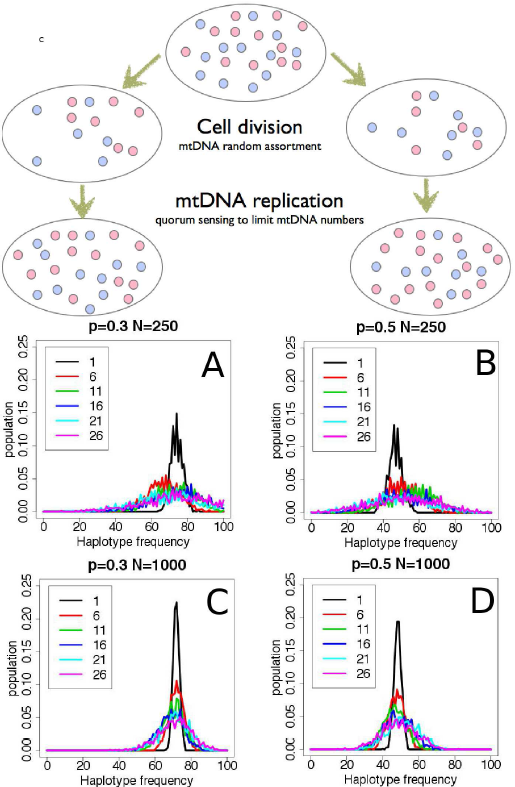
Simulations of mtDNA replication. Each cell contains a mixture of mtDNA haplotypes, here we consider only two species, red and blue, for the sake of simplicity. At cell division, the mitochondria assort randomly between the daughter cells and divide (with mtDNA replication) till a quorum of mitochondria, specific to each tissue-type, is reached. Quorum sensing is not well-understood, but the number of mitochondria per cell is tissue-specific[8]. The graphs show simulations of the evolution of heteroplasmy over time, based on this model. The distribution of haplotype frequencies spreads over time (drift), implying two randomly selected cells are unlikely to have the same mtDNA haplotype distribution. The plots show the distribution of frequencies after 1, 6, 11, 16, 21 and 26 divisions. The starting number of mitochondria per cell (N) is 250 in the upper panels (A,B) and 1000 in the lower panels (C,D). One of the alleles occurs with a frequency (p) of 0.3 in the left panels (A,C) and p=0.5 in the right panels (B,D). The drift is slower and the distributions narrower with larger N and smaller deviations of p from 0.5.

Exchange of mtDNA between cells within a population is the simplest explanation for the uniformity of heteroplasmy and its stability. Exchange can counteract the effects of drift by bringing the haplotype distribution closer to the average across the population. Other explanations, such as a balancing selection[37] or segregated replication of groups with fixed composition[38] could also be invoked to explain the lack of drift, but involve complicated mechanisms. These can also be discounted because most variants are neutral and specific to each cell line, suggesting the selection needs to be different for each cell line without an obvious selective pressure.

### Experimental Demonstration of mtDNA Exchange Between Cells

In order to test the ability of mtDNA to transfer between cells, we co-cultured cell lines with distinct mtDNA heteroplasmy signatures. Since we were not sure which kinds of cells would allow transfer and if contact between cells were important, we tested mixtures of a pair of untransformed cell lines from embryo lung fibroblasts (IMR90 and WI-38) as well as four cancer cell lines (MDA-MB-157, U2OS, A382 and HCC1806).

One cell line in each of the co-cultured pair was labelled with constitutively expressed GFP, with 10-fold more non-GFP cells than GFP. After co-culturing for
6 weeks, the GFP labelled cells were isolated (either sorting by FACS or by using Neomycin resistance) and placed in culture again for upto 2 weeks, to obtain 10 million cells, which were then prepared for mtDNA sequencing. The details are given in the methods section. There were four experiments,

1. HCC1806-GFP mixed with MDA-MB-157 (both breast cancer) (FACS)
2. U20S-GFP mixed with A382 (both cancer cells) (FACS)
3. MDA-MB-157-GFP (breast cancer) mixed with IMR90 (normal fibroblast) (Neomycin resistance)
4. WI-38-GFP mixed with IMR90 (normal fibroblast cells) (Neomycin resistance)

In two cases, (WI-38-GFP, IMR90 Fig. 5, Table 1) and (HCC1806-GFP,MDA-MB-157, Tables 2), we detected variants private to the non-GFP-labelled cell-line in the co-cultured partner cell-line, suggesting the transfer of mtDNA between the cell-lines. The private variants were transferred to varying degrees, ranging from thorough mixing to no transfer, arguing against the results arising from errors in sorting or cytoplasmic/nuclear exchange between cells. The purity of the sorted cells, based on FACS, excludes nuclear exchange as an explanation for the findings. The lack of transfer in the two pairs, 1) MDA-MB-157-GFP,IMR90 (Table ST4) and 2) U20S-GFP, A382 (Table ST5) further suggests that the transfer results are not artifacts. In all cases, the cultures were visually inspected to ensure that both cell-lines were thriving.

**Figure 5.**
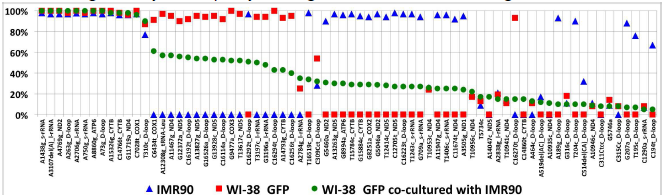
mtDNA transfer in co-cultured cells. Two normal fibroblast cell lines, WI-38 labeled with GFP and un-labeled IMR90 were co-cultured. The variants specific to each cell line before transfer and their frequency (Y-axis) is shown as red squares and blue triangles respectively. The variants are listed along the X-axis identified by the position on the mtDNA and the gene. After the co-culture, GFP negative cells were killed using the neomycin selection marker, which was incorporated with the GFP. The remaining GFP positive cells were regrown in culture and mtDNA variants sequenced (green circles). A clear transfer of variants from IMR90 to WI-38 cells is observed, as the green circles do not follow the pattern of the red squares. The variants on the X-axis are organized by the frequency of the green circles in the descending order.

### Discussion

Sensitive detection of heteroplasmy is important as its variability may have clinical significance, as a biomarker and in disease progression[39]. Analyses of heteroplasmy is confounded by Numts, highlighted by a study that used whole-genome data from the TCGA to infer that deleterious mtDNA mutations are more common in cancer cells compared to normal tissue[15]. In contrast, findings of low mutations rates in tumor mtDNA from a colorectal cancer study[14] are more in line with our findings that cancer cell lines don’t exhibit higher rates of deleterious mutations compared to normal cells from human tissues. A study of mtDNA from twins further highlights the need for further investigation of Numts[25].

Mseek provides the means to purify and deeply sequence mtDNA and determine heteroplasmy accurately by eliminating Numts and PCR-related biases. The sensitivity of Mseek increases with sequencing depth and can be made more accurate by using Duplex sequencing[33]. As currently implemented, a limitation of Mseek is its requirement of at least 4 *μ*grams of intact total DNA. Using Mseek we have demonstrated that cells from a wide-range of cell lines and human samples exhibit heteroplasmy, in accord with results from several studies[14, 19].

Heteroplasmy might be an essential feature of mtDNA, seemingly providing a *fingerprint* that can identify cells. A larger survey is needed to understand the resolution of this fingerprint and its ability to distinguish cellular origins. We found that mtDNAs from transformed human cell-lines and primary human lymphocytes are similar with respect to the distributions of densities and frequencies of mutations (benign and deleterious ones).

By studying mtDNA from colonies derived from single cells we showed that heteroplasmy is stable at the single-cell level, which is surprising in light of, 1) the higher rates of mutation in mtDNA[40] which should increase the diversity of mtDNA, and
2) drift, which should lead to homoplasmy in about 70 generations[14, 34]. Another study has determined the stability of heteroplasmy using PCR to track a particular mutation (A3243G)[38], they proposed multiplication of segregating units with fixed mixtures of haplotypes to explain this stability. Intercellular exchange of mtDNA is the simplest explanation for this stability, on the basis of our cell-line data (Fig. 3) in conjunction with simulations (Fig. 4) and co-culturing experiments (Tables 1, 2 and Fig. 5).

The transfer of mtDNA must occur through whole mitochondrial organelle transfer, as naked mtDNA in the cytoplasm would lead to an innate immune response[41], leading to cell death. To ensure that the results are not artifacts, we used two different methods to select the GFP positive cells, visually inspected over a 100 cells to make sure there was no contamination after the sorting and inspected the cells daily to make sure the colonies were growing stably and the cultures were not taken over by one cell line. The sequencing data also provides evidence that the results are not artifacts, as the transfer of variants is not uniform, some variants show a lot of transfer while others show very little or none. Additionally, certain pairs of cell lines do not show mtDNA transfer despite being co-cultured for over 6 weeks, further suggesting that the data is not an artifact and contact between cells might be needed to facilitate organelle transfer. It is possible that co-culturing cell lines that show transfer for a sufficiently long time will lead to a thorough mixing the mtDNA between the cells.

Intercellular transfer of mtDNA has been seen in other instances. Horizontal transfer of genetic material between species of yeasts has been shown[42]. Organelle transfer between cells through microtubule formation is increasingly a focus of many studies[43]. Exchange of mtDNA between mitochondria within a cell is facilitated through networks created by fusion, mediated by the mitofusins *Mfn1* and *Mfn2*[44], This intracellular exchange is essential for functional mitochondria; knocking out the fusins causes muscles to atrophy through the accumulation of deleterious mtDNA mutations[44]. *In vivo*, exchanges of mitochondria between cells have also been demonstrated in the rejuvenation of cells with damaged mitochondria by transfer of functional mitochondria from mesenchymal stem cells[45]. Rejuventation of cells containing damaged mtDNA by transfer of functional mtDNA from neighboring cells in culture has also been observed[46].

The present study is the first explicit demonstration of mtDNA transfer between healthy human cells in culture. Prior studies have shown transfer from cells with functional mtDNA into ones with non-functioning mtDNA using protein markers[45, 46]. The exchange of mtDNA between cells can explain its stability over the lifetime of an organism, inferred from the relative paucity of age-related disorders originating in somatic mtDNA mutations and over generations, inferred from the ability to identify the geographic origins of a person from the mtDNA sequence. The stability of mtDNA against deleterious mutations could be enhanced by a coupling between replication and transcription[47], ensuring the depletion of non-functional mtDNA by inefficiencies in replication.

The sequencing of mtDNA in cell-lines allows us to understand the nature of mtDNA variability and its maintenance in cell populations. Somatic mutations in mtDNA could play a role in various human disorders and in aging, especially when the transfer of mtDNA between cells is impeded. Thus, mechanisms involved in mtDNA transfer might be fruitful targets for therapeutic intervention. The transfer of functional mtDNA into diseased cells could be used to treat disorders arising from mtDNA defects. There is great value in surveying large populations in order to establish the normal range of heteroplasmy for use in GWAS studies. By making mtDNA sequencing economical, Mseek enables large-scale studies of heteroplasmy for GWAS applications and clinical monitoring of mtDNA in tissues.

## Methods

### Mseek

Our method of isolating and sequencing mtDNA, dubbed Mseek (Fig. S4), consists of the following steps, i) Total DNA is isolated from the sample, ii) The nDNA is digested using Exonuclease V, iii)The products are purified using Ampure beads to remove short fragments, iv) To test the results of the digestion, PCR primers specific to mtDNA and nDNA are used on 1*μ*l of the digested sample (Fig. 1B), v) the rest of the sample is fragmented using Covaris and end-repaired, vi) Barcoded adapters compatible with the sequencing platform are ligated to the fragments and vii) The universal adapters are used to amplify the library for loading on deep-sequencing instruments.

### Sample Processing

Total DNA was isolated from 500*μ*l of whole blood or cell lines using the Epicentre protocol (MC85200). DNA was eluted in 100*μ*l of TE buffer and checked for quality and quantity using a 1% agarose and nanodrop respectively. The eluted DNA was further heated at 70C for 30 minutes to inactivate any left over proteinase K that was introduced in the DNA isolation protocol. A first digestion was carried out by adding to the total DNA Sample (4-8*μ*g in 35*μ*l), the following, NEB4 10X Buffer ( 6 *μ*l), 10 mM ATP ATP (12 *μ*l), ExoV from NEB-M0345S (4 *μ*l) and H2O (3 *μ*l). The digests were left at 37C for 48 hours, heat inactivated at 70C for 30 minutes and purified using AMPure beads (Beckman Coulter). A second digestion was carried out by adding to the ExoV treated DNA (in 35 *μ*l) the following, NEB4 10X Buffer (6 *μ*l), 10 mM ATP (12 *μ*l), ExoV from NEB-M0345S (4*μ*l) and H2O (3 *μ*l). The digests were left at 37C for 16 hours, heat inactivated at 70C for 30 minutes and purified using AMPure beads (Beckam Coulter).

The primers listed in Tables ST1 and ST2 were used to detect presence of nuclear and mtDNA. Only if the digested DNA did not show significant PCR product for nuclear DNA, were the samples were processed for deep sequencing. In cases with nuclear DNA contamination, the digestions were repeated to improve mtDNA purification. We used Covaris for shearing and the Rapid DNA kit from Bioo Scientific for DNA library prep (5144-01).

### Cell Culture

293T (embryonic kidney cells containing the SV40 T-antigen), 501T (normal adult fibroblasts), U2OS (osteosarcoma), Saos-2 (osteosarcoma), HCC1806 (breast cancer), MDA-MB-157 (breast cancer) and A382 (glioblastoma) cells were grown in Dulbecco’s Modified Eagle’s Medium (Sigma-Aldrich, St Louis, MO, USA) supplemented with 10% fetal bovine serum (Sigma-Aldrich), 50 units/ml of penicillin/streptomycin (Gibco, Grand Island, NY, USA). All cells were sub-cultured or collected using 0.05% trypsin-EDTA (Gibco) and maintained at 37C and 90% humidity in a 5% CO2 incubator. For selection of cells carrying the neomycin selection marker, 750*μ*g/ml G418 (Gibco) was added to the culture medium for 2 weeks. Control cells not carrying the resistance marker were used to verify cell death by G418. Clonal isolation of tumor cells was performed by serial dilution into 96-well plates and visual examination of wells for single cells. Single cells were then expanded for an additional 28-30 population doublings, expanding into larger tissue culture plates as necessary.

### FACS Sorting

Cultured cells were collected in PBS (Gibco) at a density 5 × 10^6^ cells/ml and passed through a 40*μ*m filter. Cells were sorted using the BD FACS Aria II cell sorter (Becton-Dickinson, Mountain View, CA, USA), using the 488nm laser. Sorting was performed in a sterile BSL2+ biosafety cabinet. FACSDiva Version 6.1.2 software (Becton-Dickinson) was used for analysis.

### Lentiviral GFP Expression

To ectopically express GFP, we used a NSPI-derived lentiviral vector that drives GFP expression with a constitutive PGK promoter and contains a neomycin selection marker[48]. High titer lentiviral production and infection were carried out as previously described[48].

### Mixing Experiments

Two strategies were used for selecting GFP-positive cells in the mixing experiments.

In the first strategy, HCC1806 and U2OS cells were transduced with a high titer of lentivirus to constitutively express GFP. Four days after infection, GFP-positive cells were isolated by utilizing the BD FACSAria II cell and FACSDiva Version software. GFP-positive cells were placed back in culture. An examination of several hundred cells showed that all were GFP positive, suggesting that FACS sorting was highly efficient and there were no GFP-negative cells.

Pairs of cell lines (one GFP-positive and the other GFP-negative) were mixed and co-cultured in 150mm tissue culture dishes. In one group of experiments, MDA-MB-157 was co-cultured with HCC1806-GFP (ratio of 10:1 ensured by counting cells) In a second group of experiments, A382 cells were co-cultured with U2OS-GFP cells. The cells were allowed to grow for 4 weeks and were sub-cultured 1:4 or 1:5 when they reached near confluency, which was about every four days. The culture was monitored visually to ensure both GFP-positive and GFP-negative cells existed in culture.

Live GFP positive and GFP negative cells were separately gated and sterile sorted using the BD FACSAria II cell and FACSDiva software according to GFP status. Sorted GFP-positive cells were placed back into culture and allowed to expand for a week or two, to obtain 5 × 10^6^ cells in a 150 mm plate, giving ≈ 4.5 × 10^6^ of unlabelled cells and 5 × 10^5^ of GFP-labeled cells, which was then processed for mtDNA sequencing. The purity of the cells after sorting was further verified by visual examination under the microscope.

The second strategy was similar to the above with the following changes. 1) After the infection with the high titer GFP lentivirus with neomycin resistance, GFP-positive cells were selected for neomycin resistance. 2) After completion of the mixing time period, GFP negative cells were specifically selected against using the neomycin selection marker as an alternative to cell sorting. Visual examination was used to further verify the purity of the GFP positive population. In this way, only GFP positive/neomycin resistant cells were collected for mtDNA isolation. MDA-MB-157-GFP mixed with IMR90, and WI-38-GFP mixed with IMR90 were processed by this strategy.

### Analyses

The sequencing data is generated as fastq format files. These were processed using a pipeline developed by us for whole-exome analysis called MiST which is described elsewhere[35]. In brief, the sequences were filtered for quality (sequences with more than 10 consecutive nucleotides with Q < 20 were eliminated) and mapped to the reference mitochondrial genome (accession NC_012920 from Genbank). Identical reads were identified as being clonal and were considered only once, irrespective of the number of copies, towards variant calling. A variant call was made only if there were at least three non-clonal reads carrying the variant, at least 10nt away from the ends, and a minimum coverage of ten was required at the variant.

Variants occurring on reads predominantly on one strand (> 80%) of the mtDNA were excluded to further reduce errors, based on our prior experience[35]. The error rate in Miseq and Hiseq reads are usually less than 1 in a 1000 (phred score Q > 30), requiring at least 3 non-clonal reads reduces the error rate to well under 1 in a million. Nuclear contamination was estimated using sequences that map to repeat elements such as LINEs and SINEs, which only occur in nDNA. This enables reliable estimation of the level of nDNA contamination, which ranged from 0.5 – 1.5%.

The annotation of variants was determined using mtDNA annotations from MIT-OMAP13. Common SNPs (and haplotype indicators) were identified from dbSNP14. Various programs that annotate the effect of variants on protein function were tested. We eliminated programs that indicated common SNPs in mtDNA proteins were deleterious, *Mutation Assessor*[36] performed the best under this test, we used it to assess the impact of mtDNA mutations on protein function.

*Mutation Assessor* uses conservation of structure across orthologues to identify mutations in the DNA (and consequent changes in amino-acids) with potentially deleterious effects. The mutations are rated *high, medium, low*, or *neutral* based on their impact on protein function. We highlight the *high* and *medium* impact mutations in our graphs, as they might affect mitochondrial function.

Custom code was developed for simulations and the plots were created using Gnuplot and R.

## Author’s Contributions

AJ and RS designed Mseek and designed several experiments. SA and MW designed key experiments with cell-lines. LL provided human samples and suggested applications. SG and MT compared Mseek to other kits. AJ, JS, RL used Mseek on samples, EB performed cell-line work. AJ and RS wrote the paper.

## Acknowledgements

RS, AJ, LL were partially supported by the grant 1R21HG007394-01 from the NIH. Some experiments were also funded by a pilot grant from the Venture Capital Research Funding Program of Children’s Environmental Health Center (CEHC) at Mount Sinai. Brian Brown and Eirini Papapetrou helped craft the message. Comments from Viviana Simon, and Benjamin tenOever added clarity. Avinash Waghray, Sunniva Bjorklund and Vessela Kristensen caught numerous errors and gave many suggestions to improve the writing.

## Supplementary Material

Comparison of Mseek to other techniques

We tested several kits on the market using PCR with a combination of nuclear DNA and mtDNA specific primers and followed it up with sequencing only if the PCR data looked promising. PCR-based approaches (e.g. Life technologies) can be excluded for reasons discussed in the introduction. Approaches using SNP arrays (23andMe) don’t have the resolution offered by Mseek. Most other kits on the market for mtDNA purification involve lysing cells to release the organelles either chemically or by use of a dounce followed by the use of one of the following technologies,

- Purification of mitochondrial DNA from organelles using Enzyme B mix from Abcam to clean up the DNases and other proteins (kits from Abcam, PromoKine and BioVision),
- Magentic capture of mitochondria using anti-TOM22 microBeads to target the translocase of the outer mitochondrial membrane 22 (TOM 22) (MACS from Miltenyi Biotec)
- Amplification from total DNA with mtDNA specific primers (REPLI-g from Qiagen),
- Extraction of an mtDNA-enriched fraction using Qiagen’s plasmid miniprep kit followed by bead purification using Agencourt AMPure XP system[49].

We first tested the mini-prep method of isolating pure mtDNA[49]. Approximately 20 million cells (HEK 293T) were used for each isolation. The cells were loaded onto a spin mini prep column and isolation was done according to the manufacturer’s protocol (QIAGEN spin mini prep catalog # 27115). The mtDNA which is similar to plasmid DNA (size and circular structure) was eluted in 100 *μ*l of elution buffer. The mtDNA enriched fraction was later purified using the AMPure XP system. A 0.4 X proportion of beads by volume were added, collected on the magnetic stand and washed twice with 80% ethanol. Post washing, the mtDNA was resuspended in 25 ul 0.1 X TE buffer. On testing with PCR, nDNA bands were seen(Fig. S5A), consistent with the published data[49] showing that the product had about 78% nuclear DNA[49]. In that study, a subsequent mtDNA-specific amplification yielded high purity mtDNA.

We also tested the Miltenyi biotech Mitochondria MidiMACS starting kit, human (catalog # 130-094-872), since it is used by many commercial kits. Mammalian cells (≈ 10^7^) were lysed and mitochondria were magnetically labeled with Anti-Tom22 microbeads which bind to the translocase of the outer mitochondrial membrane 22 protein (TOM 22). The sample was then loaded onto the column placed in the MACS separator. After washing only magnetically labeled mitochondria are retained on the column. The column is detached from the separator and mitochondrial organelles are eluted. Mitochondrial DNA is further isolated from the organelles by isopropanol precipitation. Since nuclear DNA remained in the MACS-purified samples according to PCR(Fig. S5B), consistent with the published data[49], sequencing was not performed.

We compared Mseek to REPLI-g (Qiagen), using sequencing on mtDNA from a blood sample (in triplicate) to establish that Mseek exhibits the highest purity of mtDNA compared to kits currently available on the market(Fig. S5C). Mseek data from cell lines shows substantial reduction of nuclear DNA and uniform coverage across the length of mtDNA(Fig. 1).

**Figure S1.**
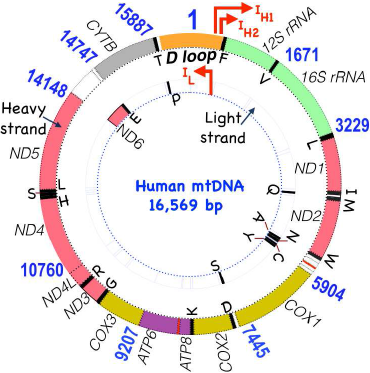
Organization of mtDNA. 13 protein-coding genes, 22 tRNA genes and 2 rRNA genes are encoded by a single circular nucleic acid and transcribed from three promoters: LSP (inner circle of genes - strand), HSP1 (outer circle of gene, + strand) and HSP2 (16S rRNA) on the D-loop, which is non-coding but critical for replication and transcription. The three polycistronic transcripts are processed by enzymatic excision of the tRNAs. There are a few small gaps (< 30 nt) in annotation, and a 45nt overlap between ATP6 and ATP8 which might have roles in replication. The mitochondrial genetic code differs from the nuclear code. In mammalian mitochondria, *ATA* codes for Methionine instead of Isoleucine, *TGA* codes for Tryptophan instead of the stop codon, and *AGA, AGG* code for stop codons instead of Arginine hinting at a bacterial origin[50].

**Figure S2.**
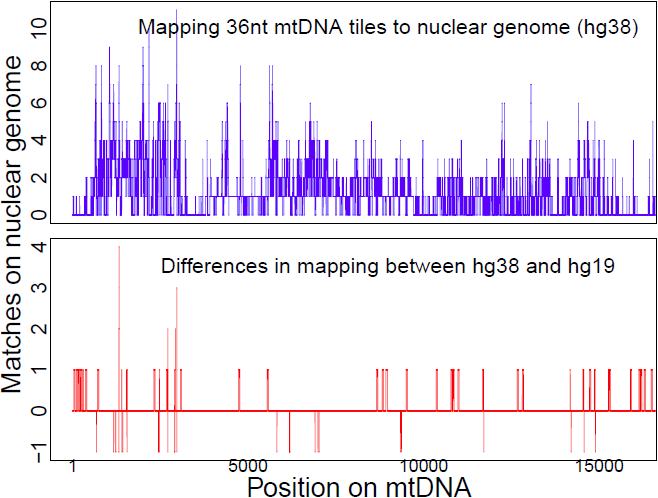
Numts (Nuclear-mtDNA pseudogenes). The x-axis is the position along the mtDNA. The graph on the top shows the number of matches on the nuclear genome (hg38) of 36 nt tiles from the mtDNA. The graph in the bottom shows the changes in the mapping numbers for the tiles between hg38 and hg19, positive numbers are an increase in matches, while negative numbers are a decrease.

**Figure S3.**
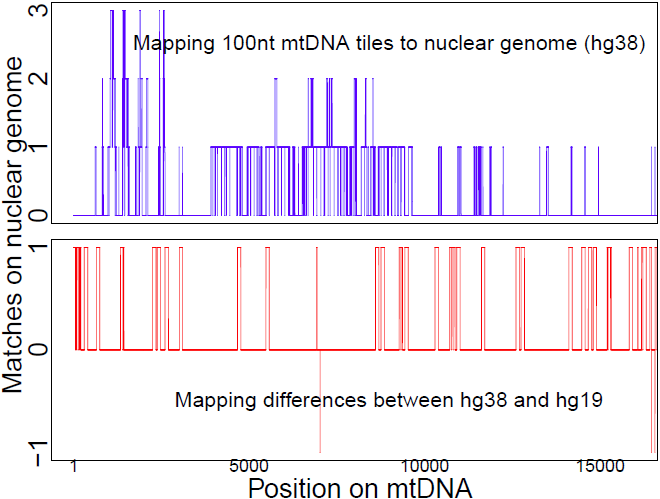
Numts (Nuclear-mtDNA pseudogenes). The x-axis is the position along the mtDNA. The graph on the top shows the number of matches on the nuclear genome (hg38) of 100nt tiles from the mtDNA. The graph in the bottom shows the changes in the mapping numbers for the tiles between hg38 and hg19, positive numbers are an increase in matches, while negative numbers are a decrease. Even at 100nt, there are a number of Numts in the reference genome.

**Figure S4.**
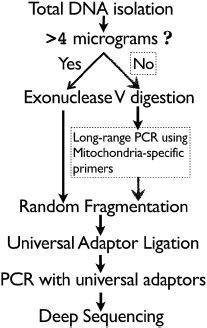
The Mseek protocol.

**Figure S5.**
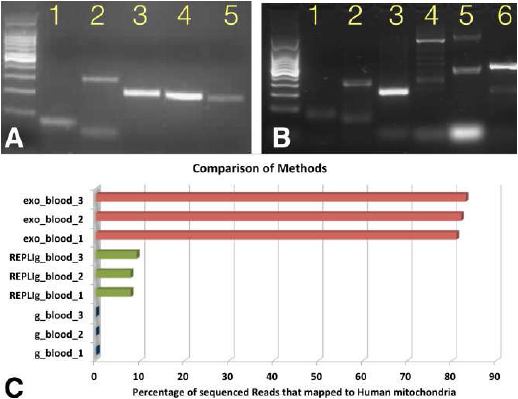
**Mseek comparisons to other techniques**. Plasmid mini-prep (panel A) and MACS (panel B) show a lack of purification of mtDNA based on PCR products run on a gel, thus they were not prepared for sequencing. Panel C shows sequencing data from REPLI-g (Qiagen) compared to Mseek, both of which show purification of mtDNA on the gel. A Plasmid mini prep followed by bead purification. PCR using primers specific to 18S(1), 28S(2), B-actin(3), mtDNA(4), mtDNA(5) show nDNA is present after treatment. B MACS. PCR using primers specific to 18S(1), 28S(2), B-actin(3), GAPDH(4), mtDNA(5), mtDNA(6) show that nDNA is present after prep.C Deep-sequencing data. Mseek (red bars) exhibits consistently better enrichment of mtDNA compared to REPLI-g (green bars) and untreated total DNA (blue bars). This comparison was done in triplicates, each using 8 *μ*g of total DNA from the same blood sample. The purity of mtDNA from Mseek (80%) is lower compared to results presented in the text from cell lines, due to various factors in blood which requires further fine-tuning.

**Table S1.**
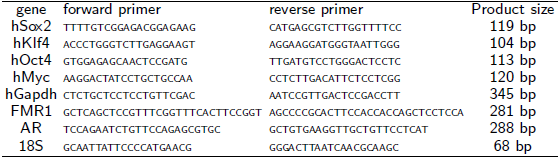
Primers specific to human nuclear DNA.

**Table S2.**
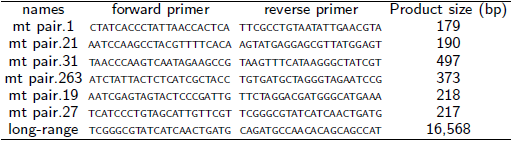
Primers specific to human mtDNA. The last row is the primer pair for amplifying the whole mtDNA using long-range PCR.

**Table S3.** Transitions versus Transversions for data from IMR90, WI-38 and MDA-MB-157. Of a total of 84 single base mutations there are only 6 transversions, the rest are transitions. The row labels are the original reference base, the column labels are the variant bases, so G to A changes happened 15 times. Most mutations were transitions (Table ST3), suggesting that the mutations don’t arise from oxidative stress, and are most likely driven by the errors introduced by polymerase-γ activity[33].

|  | a | c | g | t |
| --- | --- | --- | --- | --- |
| A | 0 | 1 | 23 | 0 |
| C | 2 | 0 | 0 | 17 |
| G | 15 | 3 | 0 | 0 |
| T | 0 | 23 | 0 | 0 |

**Table S4.**
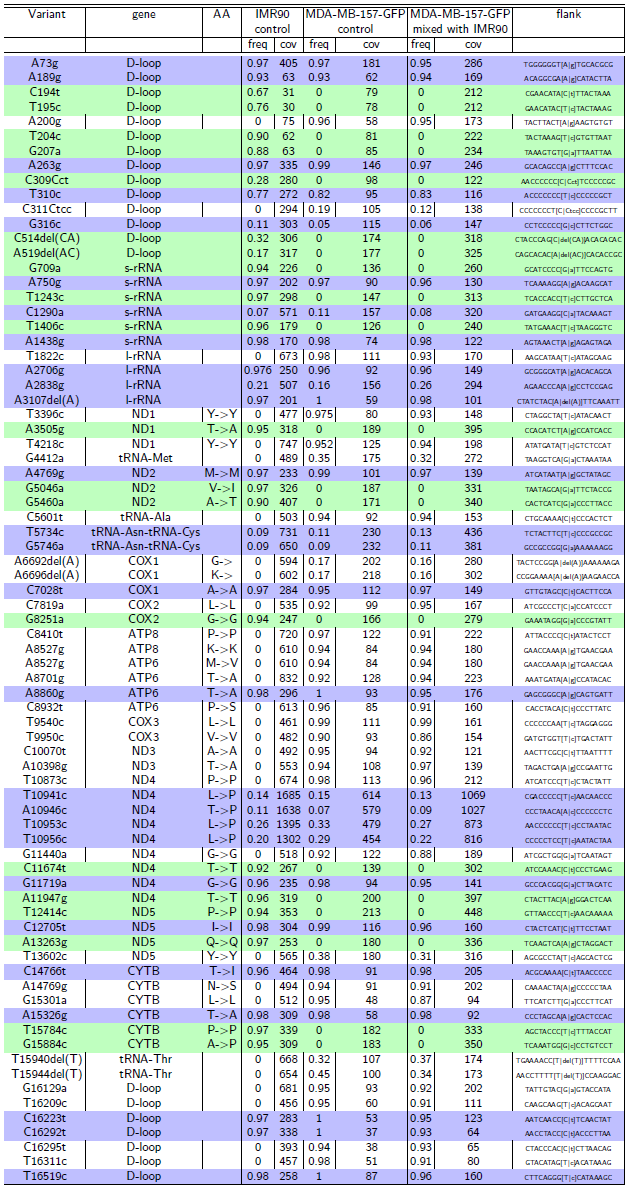
Co-culturing experiments for MDA-MB-157-GFP(cancer) with IMR90(fibroblast) The white rows are variants private to MDA-MB-157-GFP, gray rows are variants that transferred from IMR90 to MDA-MB-157-GFP and blue rows are variants common to IMR90 and MDA-MB-157-GFP. There were no transfers in this case. The frequency is *freq* (which ranges from 0-1) and the coverage, number of reads covering the variant, is *cov*.

**Table S5.**
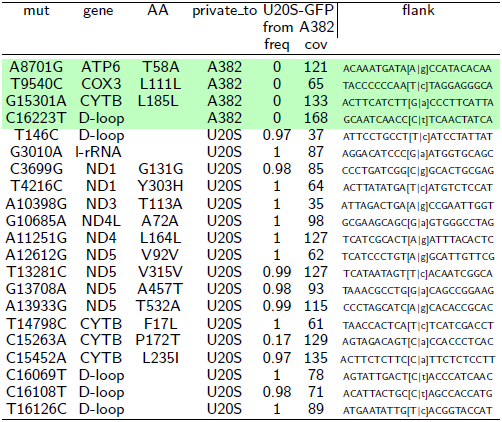
Data from mixing experiments of U20S-GFP co-cultured with A382 for 6 weeks. The column private_to identifies the cell-lines that exhibit the variant. The light green rows are variants private to A382 that did not transfer into U20S. The frequency is *freq* (which ranges from 0-1) and the coverage, number of reads covering the variant, is *cov*.

